# The effect of cold exposure on energy expenditure of mice fed an obesogenic diet

**DOI:** 10.1101/2025.09.29.679278

**Authors:** Angelica Amorim Amato, Andreza Fabro de Bem, Richard Cheng-An Chang, Riann Egusquiza, Bruce Blumberg

## Abstract

**Objective:** Cold exposure is one of the most powerful physiological stimuli for thermogenic adipose tissue activity and may positively impact metabolic homeostasis. An important gap in knowledge is whether mice with diet-induced obesity respond to cold similarly to their lean counterparts. The goal of the present study was to compare the response to cold exposure between mice fed a standard (normal-fat) diet and a higher-fat diet.

**Methods:** Male C56BL/6J mice fed a standard diet (13.1% fat) or a higher-fat diet (21.6% fat) during adulthood and exposed to cold (4-6°C) following different approaches. Body weight, body composition, food intake, rectal temperature, energy expenditure, and respiratory exchange ratio were assessed.

**Results:** Cold exposure significantly increased energy expenditure and limited weight gain despite elevated food intake in mice fed either a standard or a higher-fat diet, while body composition, body temperature, and respiratory exchange ratio remained stable across both dietary groups. Moreover, energy expenditure measured at 4-6 °C was comparable between mice fed standard and higher-fat diets, demonstrating that cold-induced thermogenesis may elicit a consistent metabolic response independent of dietary fat content.

**Conclusions:** Our results demonstrated that cold exposure increased energy expenditure in both lean and obese animals, highlighting thermogenesis as a promising target for obesity treatment beyond current approaches focusing on appetite suppression.

**Highlights:**

- Cold exposure increases energy expenditure and limits weight gain in both control diet and higher-fat diet-fed mice, despite increased food intake.
- Core body temperature and body composition remain stable during cold exposure, regardless of dietary fat content.
- Cold-induced thermogenesis elicits a comparable metabolic response in control diet and higher-fat diet-fed mice, supporting its potential as a therapeutic strategy for obesity.

## 1. Introduction

In mammals, thermogenic adipose tissue is specialized for the oxidation of nutrient substrates to generate heat through non-shivering thermogenesis. This process is primarily driven by the action of uncoupling protein 1 (UCP1), although UCP1-independent thermogenic mechanisms, such as creatinine-substrate cycling ^1^ and calcium cycling through the sarco-endoplasmic reticulum ATPase 2b ^2^, have also been described.

Two types of thermogenic cells, brown and beige adipocytes, are present in both rodents and humans. Brown adipocytes arise from the dermatomyotome and reside in dedicated depots in rodents and human infants. In contrast, beige adipocytes represent a heterogeneous and inducible population of thermogenic cells located within white adipose tissue. In the adult human, thermogenic fat is predominantly located in the supraclavicular and thoracic regions ^3^, where UCP1-expressing adipocytes exhibit a molecular signature closely resembling that of rodent beige cells ^4^.

Beyond its established role in maintaining body temperature during cold exposure, thermogenic adipose tissue plays a critical role in regulating systemic energy balance in mice ^5^. These functions appear to extend to human adults, with evidence indicating that brown fat activation occurs during cold-induced thermogenesis ^6^. Moreover, the presence of thermogenic adipose depots is positively correlated with markers of metabolic health ^5^ and is independently linked to a reduced risk for metabolic diseases, such as type 2 diabetes, dyslipidemia, and hypertension ^7^. The later findings have increased the interest in exploring thermogenic adipose tissue as a potential target for treating metabolic diseases.

Cold exposure is the most powerful physiologic stimuli for activating thermogenic adipocytes ^8^, acting through norepinephrine/μ3-adrenergic signaling to enhance the expression of thermogenic genes and the proliferation and differentiation of thermogenic precursor cells. While the effect of cold exposure to increase energy expenditure and potentially reduce body mass has been well documented, little is known about the interplay between dietary factors and thermogenic modulation of metabolic outcomes, highlighting the need for further investigation. To address this gap, we examined the effect of cold exposure on metabolic responses in mice fed either a normal-fat diet or an obesogenic diet.

## 2. Material and methods

### 2.1. Animal maintenance

C57BL/6J mice were acquired from the Jackson Laboratory (Sacramento, CA) and housed in micro-isolator cages with 4 mice/cage. under controlled environmental conditions. Animals were kept at 25°C, prior to temperature manipulations, and under a 12 h light/dark cycle. Water and food were provided ad libitum; animals were maintained on a normal-fat/standard diet (SD-Rodent Diet 20 5053^*^, 13.1% kcal from fat; PicoLab) or fed a higher-fat diet (HFD-Mouse diet 20 5058^*^; 21.6% kcal from fat; PicoLab). All procedures were approved by the Institutional Animal Care and Use Committee of the University of California, Irvine and conducted in accordance with Federal guidelines. Measures were taken to ensure humane treatment and minimize suffering throughout the study.

### 2.2. Cold exposure

Cold exposure was conducted using three approaches. In the first approach (Figure 1A), mice fed the standard diet (n = 4) were singly housed at 10.5 weeks of age and exposed to a gradual temperature drop from 30°C to 4-6°C over 9 days, followed by a return to 25°C. Energy expenditure and respiratory exchange ratio were assessed throughout temperature decrease.

**Figure 1.**
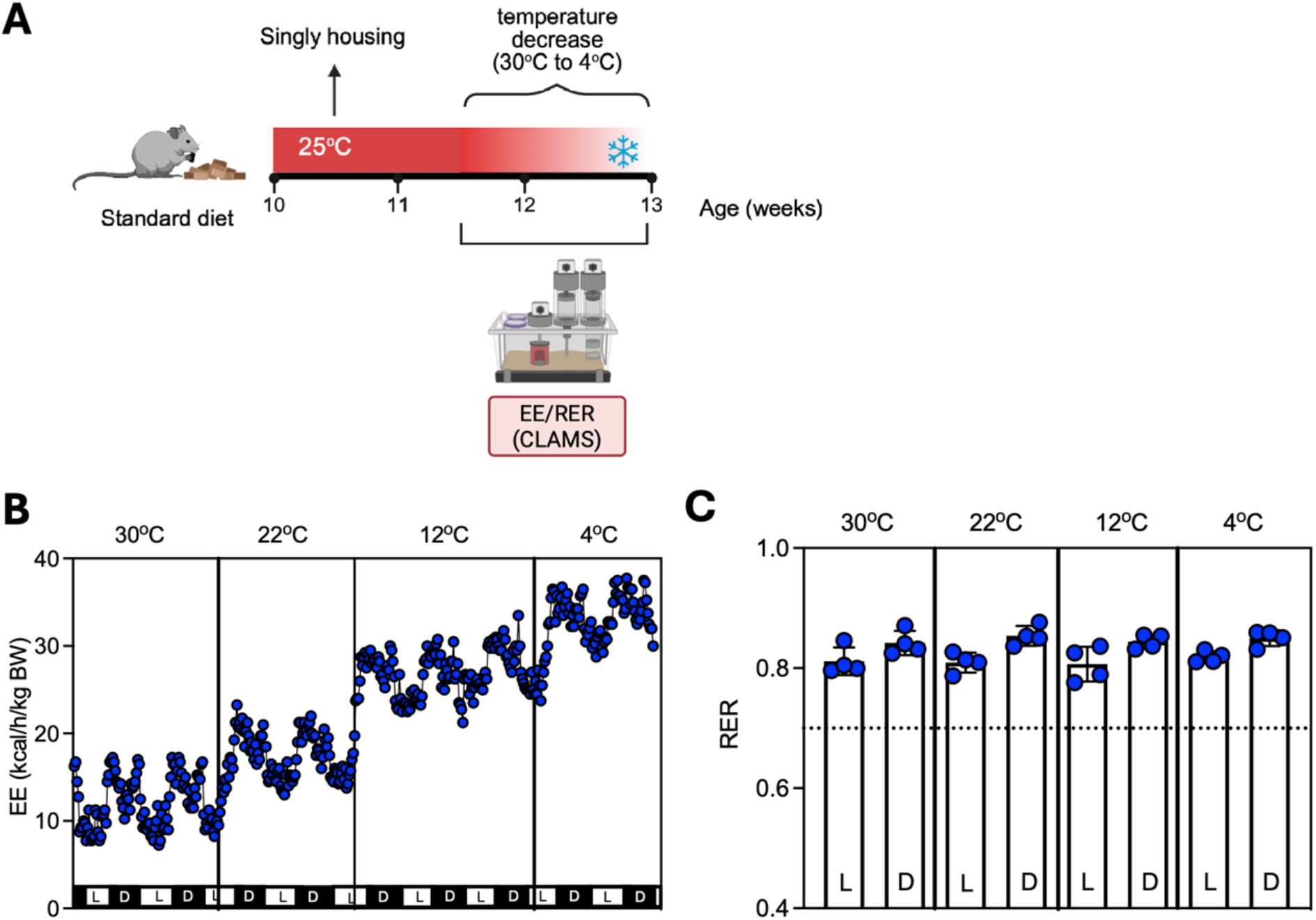
Gradual reduction in ambient temperature increases energy expenditure. (A) Mice were singly housed at 10.5 weeks of age and randomly assigned to a gradual temperature decrease from 30 °C to 4–6 °C over 9 days and assessed for (B) energy expenditure and (C) respiratory exchange ratio. N = 4 mice. CLAMS: comprehensive laboratory animal monitoring system, EE: energy expenditure, RER: respiratory exchange rate.

In the second approach (Figure 2A), ten mice, initially maintained at 25°C and fed the standard diet, were singly housed at 15.5 weeks of age. At week 16, they were placed in a cold room (4-6°C) for two weeks. Body weight was assessed every one to two days. Body composition was assessed using EchoMRI™ Whole body Composition Analyzer. Food consumption was assessed before and throughout the cold exposure period.

**Figure 2.**
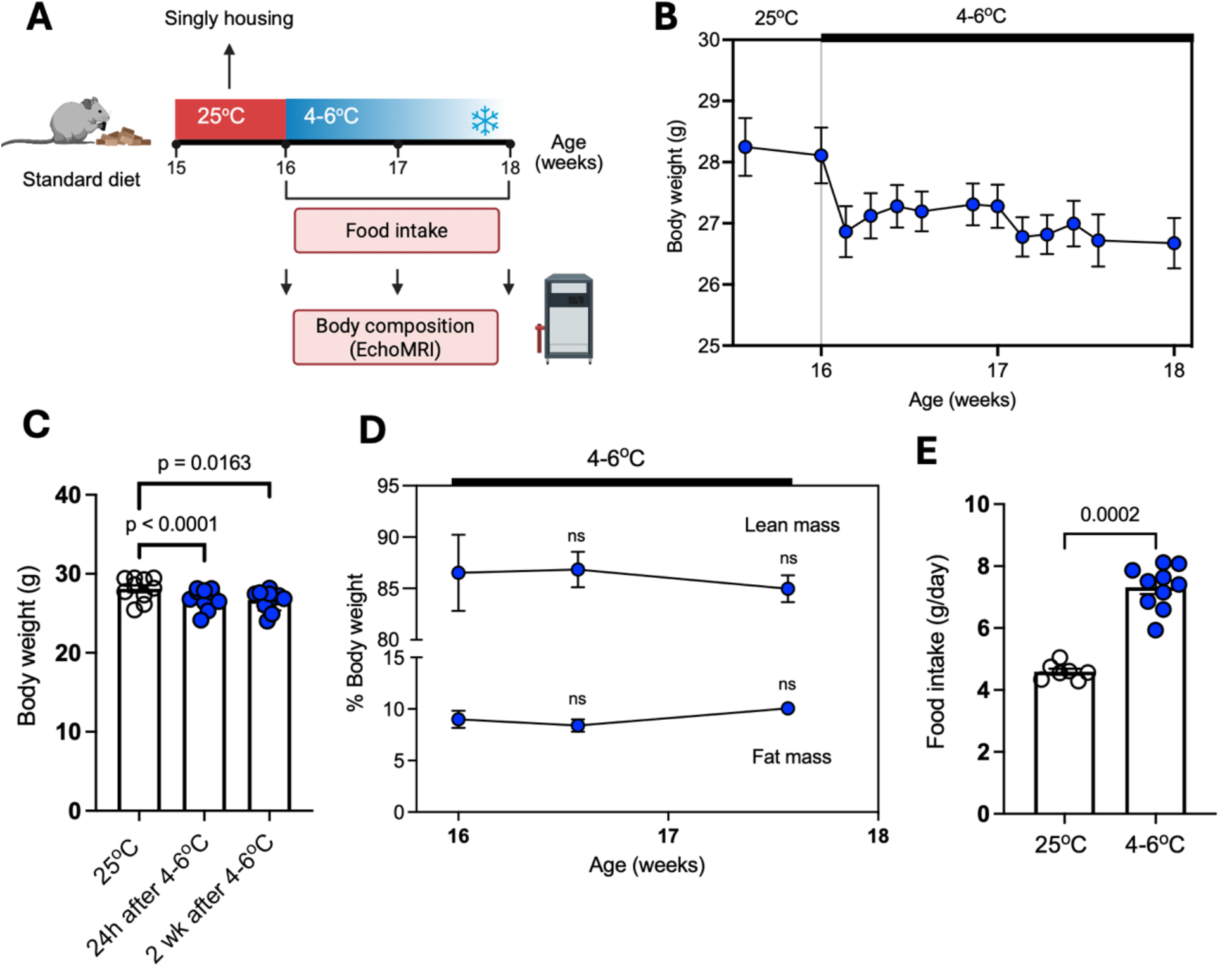
Prolonged cold exposure induces modest weight loss in standard diet-fed male mice, despite increased food intake and unchanged body composition. (A) Experimental design: mice were maintained at 25 °C, singly housed at 15.5 weeks of age, and exposed to 4–6 °C for two weeks. Assessments included (B) body weight, (C) weight gain, (D) body composition, and (E) food intake. N = 10; data in (C) and (E) were analyzed by paired t test, and in D were analyzed using two-way ANOVA.

In the third approach (Figure 3A), a subgroup of mice (n = 15, cold exposure mice) was maintained on a standard diet until 13 weeks of age, after which they were fed the higher-fat diet. These mice were singly housed at 13 weeks and placed in a cold room (4-6°C) at 15 weeks of age, for four weeks. Rectal temperature was measured four hours after the onset of cold exposure, and then daily, using a rectal probe (RET-4, Physitemp). Body weight was assessed daily throughout the cold exposure period. Food consumption was assessed weekly before and two weeks after the onset of cold exposure, by calculating the difference between diet mass in the hopper over seven consecutive days. During the final week of cold exposure, energy expenditure and respiratory exchange ratio were assessed at 4-6°C for 2.5 days. A second subgroup of mice (n = 8, control mice) was maintained on a standard diet until 15 weeks of age, after which they were singly housed and switched to a higher-fat diet. The latter group remained at 25°C throughout the entire experimental protocol, with body weight and food consumption assessed weekly. Both subgroups underwent body composition assessment weekly, using EchoMRI™ Whole body Composition Analyzer.

**Figure 3.**
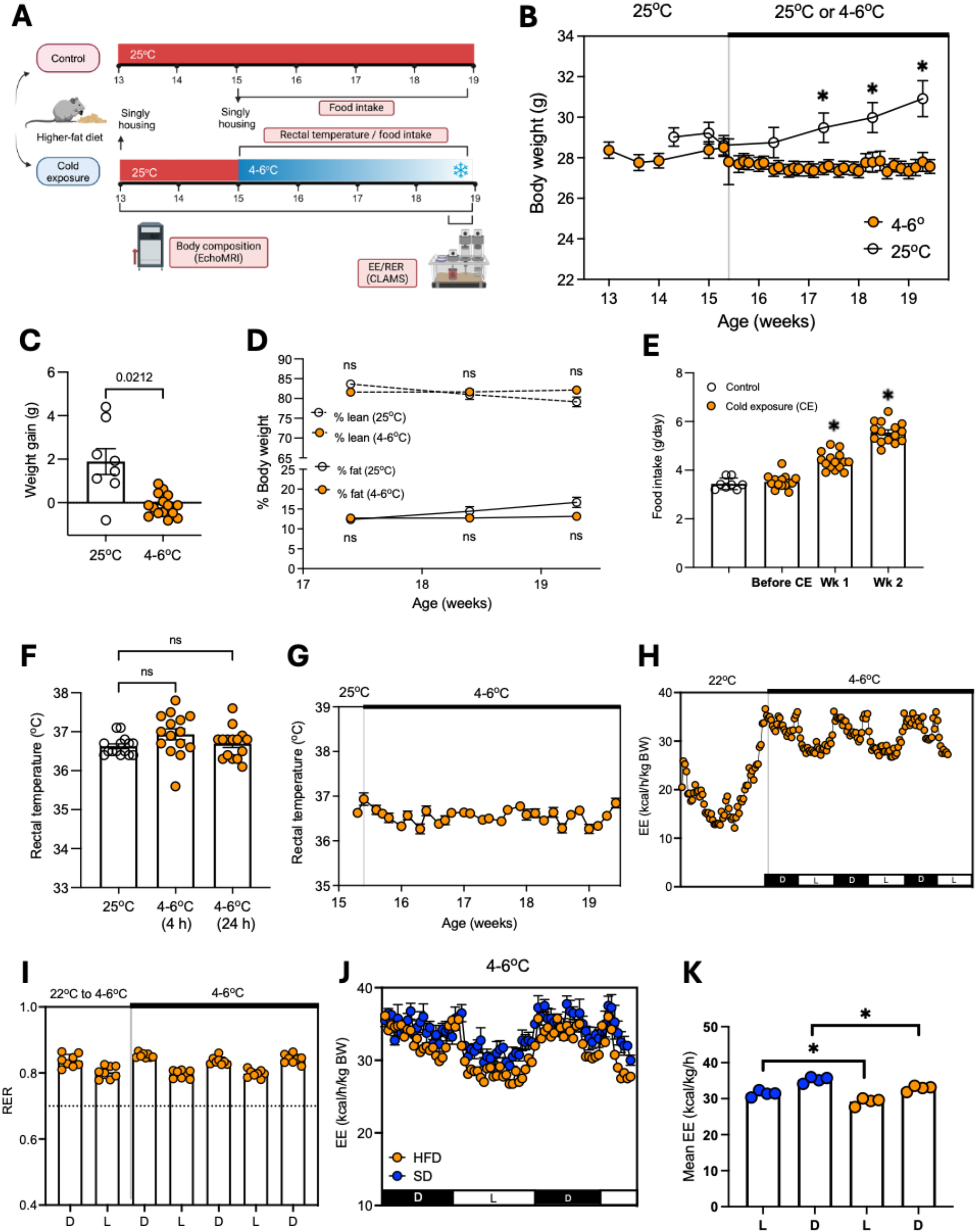
Prolonged cold exposure attenuates higher-fat diet-induced weight gain and adiposity while increasing energy expenditure and food intake in male mice. (A) Experimental design: mice maintained at 25 °C were singly housed and randomly assigned to cold exposure (4-6 °C, n = 15) for four weeks or to remain at 25 °C (n = 8). Assessments included (B) body weight, (C) weight gain, (D) body composition, (E) food intake, (F) rectal temperature before and after 4 and 24 h of cold exposure, (G) rectal temperature throughout the cold exposure period, (H) energy expenditure (EE), and (I) respiratory exchange ratio (RER). (J, K) Energy expenditure at 4-6 °C in mice fed the standard diet and gradually exposed to temperature decrease over 9 days versus those fed a higher-fat diet-fed exposed to 4-6 °C for four weeks. Data in (C), (E), (F), and (K) were analyzed by paired t test, and in B and D were analyzed using two-way ANOVA. ^*^ p < 0.05 (vs control in E). CE: cold exposure.

### 2.3. Indirect calorimetry

Oxygen consumption (VO_2_), carbon dioxide production (VCO_2_), and respiratory exchange ratio (RER, VCO_2_/ VO_2_) were determined using the TSE PhenoMaster® System (TSE Systems, Homburg, Germany). Energy expenditure was calculated as the product of the caloric value of oxygen (3.815+1.232 x RER) and VO_2_ and expressed as kcal/kg body weight/h.

## 3. Results

### 3.1. The effects of cold exposure in standard diet-fed mice

To investigate the effects of cold exposure, standard (normal-fat) diet-fed mice housed at 25°C were subjected to one of two temperature protocols: ambient temperature was either gradually reduced to 4-6°C following an initial increase to thermoneutrality (30°C; Figure 1A), or directly decreased to 4-6 °C for two weeks (Figure 2A). In the first protocol (Figure 1A), the transition from 30 °C to 4-6 °C resulted in a substantial increase in energy expenditure - approximately threefold during the light cycle and twofold during the dark cycle (Figure 1B) - without affecting the respiratory exchange ratio (RER) (Figure 1C).

In the second protocol (Figure 2A), over the 14-day cold exposure period, mice exhibited a modest but significant weight loss of 5.1% relative to baseline, beginning at the first day of temperature decrease (Figures 1B-C). Body composition did not change significantly (Fig. 2D), whereas food intake increased ∼1.8-fold in response to cold (Figure 2E).

### 3.2. Effects of cold exposure in higher-fat diet-fed mice

We investigated the effect of cold exposure (4-6°C) versus ambient temperature of 25°C over a four-week period in mice fed a higher-fat (Figure 3A). Mice were singly housed 24 hours prior to cold exposure, which induced a slight weight decrease. However, body weight remained stable throughout the entire cold exposure period (Figure 3B). In contrast, mice maintained at 25°C and fed the same diet showed a progressive increase in body weight (Figure 3B). Therefore, there was a significant difference of weight gain in mice maintained at each temperature (Figure 3C). While body composition remained stable in the cold-exposed mice, those kept at 25°C exhibited an increase in body fat percentage (Figure 3D).

Food consumption significantly increased after exposure of mice to 4-6°C (Figure 3E). There was no significant change in rectal temperature before (25°C) or following the first hours after temperature decrease (4-6°C), although one mouse exhibited rectal temperature decrease to less than 36°C shortly after being exposed to cold (Figure 3F). Moreover, mean rectal temperature remained stable and above 36°C throughout cold exposure (Figure 3G). When mice were removed from the cold room and placed in the calorimeter, we observed an increase in energy expenditure following temperature decrease from 22°C to 4-6°C (Figure 3H), while the respiratory exchange ratio remained similar before and after temperature reduction (Figure 3I).

We directly compared the energy expenditure at 4-6 °C between mice fed with each diet. Despite differences in age (11 vs. 18 weeks) and cold exposure protocols (gradual decrease from 30 °C to 4–6 °C over 9 days versus sustained exposure to 4–6 °C for three weeks) between standard and higher-fat diet-fed mice, we found that energy expenditure of mice fed a higher-fat increased in response to temperature decrease and was only slightly lower than that of mice fed a standard diet (Figure 3J and 3K). These differences were 29.2 ± 1.1 at 4-6°C vs 31.4 ± 0.8 kcal/kg/h at 25°C during the light phase and 32.9 ± 0.8 at 4-6°C vs 35.2 ± 0.8 kcal/kg/h at 25°C during the dark phase.

## 4. Discussion

This study showed that male C57BL/6J mice fed an obesogenic diet exhibited increased energy expenditure in response to cold exposure, comparable to mice maintained on a standard normal-fat diet. This response was characterized by an initial reduction in body weight, followed by stabilization of both body weight and fat mass during prolonged cold exposure, likely driven by compensatory increases in energy intake. These findings indicated that cold exposure can mitigate weight gain and adiposity typically induced by higher-fat feeding, underscoring the capacity of cold exposure to modulate metabolic outcomes even in the context of an obesogenic diet.

Cold exposure induces a rise in energy expenditure in mammals to maintain core body temperature primarily through two pathways: shivering thermogenesis in skeletal muscle and non-shivering thermogenesis in thermogenic adipose tissue ^5^. However, the effects of cold exposure on body weight and adiposity are modulated by several factors, including the intensity and duration of cold exposure, food availability, and physiological adaptations that enhance both the ability to harvest energy from food and the activation thermogenic pathways.

Healthy mice exposed to cold typically maintain their body weight ^9^ or experience a slight weight loss ^10^ during the initial 10 to 14 days. However, in animals with unrestricted access to food, this initial phase is often followed by weight gain, driven primarily by increased food intake and gut microbiota-mediated expansion of intestinal size and absorptive capacity ^9^. These observations highlight the critical influence of food availability and the body’s ability to efficiently harvest energy from the diet in regulating body weight and adiposity during cold exposure. While gut microbiota-induced intestinal remodeling has been implicated in enhanced energy harvesting, the molecular mechanisms that couple elevated energy intake with increased energy expenditure remain poorly understood and require further investigation.

Mice exposed to cold for 14 days exhibited elevated circulating leptin levels, paralleling the decreased fat mass during this period ^10^. However, increased leptin alone is unlikely to account for the sustained increase in food intake observed during prolonged cold exposure, particularly when body fat was shown to increase ^9^. The involvement of other humoral factors in regulating appetite, including thermogenic adipose tissue-derived batokines, remains largely unexplored. Notably, increased food intake has also been observed in humans exposed to mild cold, even in the absence of measurable increases in energy expenditure ^11^. These findings raise the possibility that cold exposure may directly promote feeding behavior, an effect that warrants further mechanistic investigation.

Although the mechanisms by which cold exposure influences appetite and satiety remain incompletely understood, our findings, along with those from previous studies ^9,10^, suggest that reductions in body weight and adiposity during cold exposure likely require some degree of caloric restriction. Both standard diet and higher-fat diet-fed mice increased their food intake at 4-6°C, yet body weight and fat mass percentage remained stable. In higher-fat diet-fed mice, cold exposure mitigated the diet-induced weight and fat gain, despite not inducing net loss in either parameter. These results indicated that the increased energy demands imposed by cold are largely compensated by increased caloric intake, and that a negative energy balance achieved through food restriction is likely necessary for weight loss under these conditions. Future studies should assess whether thermogenic responses can be sustained under caloric deficit, and to what extent energy expenditure adapts to restricted energy availability during cold exposure.

A central question in addressing the therapeutic potential of thermogenic adipose tissue activation for obesity is whether obesity itself impairs the tissue’s responsiveness to thermogenic stimuli, in both rodents ^12,13^ and humans ^14^. This raises the concern that cold exposure may be less effective at improving obesity and related metabolic dysfunction once these conditions are established. Previous rodent studies have shown that diet-induced obesity led to whitening of brown adipose tissue, characterized by reduced expression of thermogenic markers ^12^ impaired responsiveness to β3-adrenergic signaling ^13^ in brown adipose tissue, diminished energy expenditure ^12^, and lower core body temperature ^12^.

In contrast, our findings indicated that mice fed a higher-fat diet retained the ability to increase energy expenditure in response to cold (4-6°C), comparable to mice on a standard diet. This discrepancy may reflect differences in dietary composition between studies; for example, the fat content in our study (21.6%) was substantially lower than in prior work (typically 50-60%), which may differentially affect brown adipose tissue function. Additionally, it remains possible that impairments of thermogenic capacity at higher temperatures do not preclude recruitment of thermogenesis under more extreme stimuli such as that assessed herein. These findings suggest that thermogenic responses can be preserved under certain obesogenic conditions, although further studies are needed to determine whether this holds true in the context of more severe dietary challenges, including high-fat, high-sucrose, or combined obesogenic diets.

## 5. Conclusions

Our findings indicated that cold exposure enhanced energy expenditure in both lean and obese mice, highlighting the potential of thermogenic activation as a strategy to promote negative energy balance. These results underscore the therapeutic relevance of targeting thermogenesis for obesity treatment. Given that most current interventions primarily aim to reduce caloric intake, our data suggested that increasing energy expenditure, whether through cold exposure or by pharmacologically mimicking its effects, may offer a complementary or alternative approach for long-term weight management.

## Funding

Supported by a grant from the US Public Health Services (NIH R01 ES031139) to BB and from the Foundation for Research Support of the Federal District, Brazil (00193-00002154/2023-84) to AAA.

## Declaration of interest

none.

## CRediT author contribution statement

Angélica Amorim Amato: Conceptualization, Data curation, Formal analysis, Investigation, Methodology, Supervision, Visualization, Writing – original draft.

Andreza Fabro de Bem: Data Curation, Formal analysis, Writing – original draft.

Richard Cheng-An Chang: Formal analysis, Investigation, Methodology, Writing – review & editing.

Riann Egusquiza: Formal analysis, Investigation, Methodology.

Bruce Blumberg: Conceptualization, Funding Acquisition, Methodology, Resources, Supervision, Writing – review & editing.

## Notes

### Competing Interest Statement

The authors have declared no competing interest.

